# Mechanical and signaling mechanisms that guide pre-implantation embryo movement

**DOI:** 10.1101/2020.04.20.051094

**Authors:** Diana Flores, Manoj Madhavan, Savannah Wright, Ripla Arora

**Affiliations:** Department of Obstetrics, Gynecology and Reproductive Biology, Michigan State University; Department of Biomedical Engineering, Michigan State University; Institute for Quantitative Health Science and Engineering, Michigan State University

**Keywords:** Embryo spacing, Embryo-uterine interactions, Implantation, Muscle contraction, LPAR3

## Abstract

How a mammalian embryo determines and arrives at its site of attachment is a mystery that has puzzled researchers for decades. Additionally, in multiparous species, embryos face a unique challenge of achieving adequate spacing to avoid competition for maternal resources. Using our enhanced confocal imaging and 3D image reconstruction technology, we evaluate murine embryo location in the uterus along the longitudinal oviductal-cervical axis. Our analysis reveals three distinct pre-implantation stages: a) Embryo entry; b) Unidirectional movement of embryo clusters; and c) Bidirectional scattering and spacing of embryos. We show that unidirectional movement of embryo clusters is facilitated by a mechanical stimulus of the embryo as a physical object and is regulated by adrenergic uterine smooth muscle contractions. Embryo scattering, on the other hand, relies on embryo-uterine communication reliant on the LPAR3 signaling pathway and is independent of adrenergic muscle contractions. We propose that the presence of embryo clusters in the uterine horn provides an opportunity for the uterus to sense and count the embryos, followed by scattering and spacing these embryos along the given length of the horn. Thus, uterine implantation sites in mice are neither random nor predetermined but are guided by the number of embryos entering the uterine lumen. These studies have implications for understanding how embryo-uterine communication is key to determining an optimal implantation site, which is necessary for the success of a pregnancy.

**Significance Statement:** In mammals that carry multiple offspring in one gestation, embryos seemingly acquire even embryo spacing. Such even distribution would imply a guided interaction between the mother and the fetus very early on in pregnancy to allow favorable pregnancy outcomes. Thus, it is essential to understand quantitatively if and when such a uniform distribution of embryos is established. Further, uncovering the physical and biological mechanisms that allow for such equal distribution of embryos, will improve our understanding of early pregnancy events and provide for novel targets for improving pregnancy success in case of infertility and artificial reproductive technologies as well as to develop non-hormonal therapies for contraception.

## Introduction

In understanding the biology of maternal-fetal interactions, one of the crucial steps is the communication of the early embryo with the uterine milieu to find a ‘good’ site for attachment. In multiparous species, there is an additional challenge to address even spacing amongst embryos to avoid competition for maternal resources. The mouse model serves as an excellent model system to address both these questions in early mammalian pregnancy. Reproducible patterns of implantation such as attachment near the fundus in humans that are monotocous (single offspring) species (1), and even distribution of embryos along the uterine length in rats and rabbits that are polytocous (multiple offspring) species (2, 3) suggest evolutionary mechanisms that have been selected to allow for these patterns.

In 1956, Boving suggested that a detailed analysis of embryo distribution at the time of implantation reflects whether the embryo location is acquired randomly due to diffusion and muscle contraction movements or due to a pre-determined stimulator-effector system (2). Quantitative analysis of embryo spacing has been performed in the rabbit model system (2) as well as in the rat (3), and when compared to a random model, suggests non-random distribution of embryos implicating embryo-uterine, or/and embryo-embryo interactions in the definitive determination of implantation sites.

In the rabbit, embryo distribution along the uterus was measured using two features: the distance of an individual blastocyst away from the oviductal-uterine junction and the relative distance between successive embryos. Embryo-embryo distance successively increased over time, suggesting that embryos enter and move unidirectionally while separating and achieve equal spacing only before implantation. This study did note that the ends of the uterine horn, i.e., regions of the uterus closest to the oviduct and the cervix, can be refractory to the presence of embryos (2). In the rat, the distribution of embryos was analyzed after entry into the uterus until the time of implantation. For each time point, the uterus was cut in three segments of equal length, and the number of embryos located in each segment was quantified by flushing these segments. All embryos are initially present in the rostral segment – closer to the oviduct. Over time, the embryos seem to distribute in the first and second segments and, eventually, in all three segments equally (4). Such distribution of embryos, suggests that in the rat, similar to the rabbit (2), embryos enter the uterine horn and move in a unidirectional manner, spacing out eventually before implantation. In the mouse, Restall and Bindon performed 2D histological analysis of embryo location, finding that embryos accumulated in the center of the horn before starting a bidirectional movement around 10:00h on day 3 of pregnancy (5). While they evaluated patterns of embryo movement, there was no determination of the randomness of the eventual implantation site.

Embryo movement in rabbits, rats, and mice has been attributed to uterine muscle contractions. In rabbits, spontaneous uterine contractions are responsible for moving the embryo in the preimplantation stages, by a process similar to agitation (2, 6). Contractions are also implicated in even embryo spacing, and these contractions are thought to be myogenic and not neurogenic as the denervated uterus retains the capability to contract (7). In rats, when relaxin, an inhibitor of alpha-adrenergic signaling is used, embryo movement is slowed down, leading to the accumulation of embryos in the rostral segment as opposed to the distribution of embryos along the uterine horn (4). However, once relaxin exposure is removed, the embryos are able to space out evenly for implantation (4). If relaxin is administered continuously from the time of embryo entry until the time of implantation, then overcrowding of embryos in the first third of the horn is observed at the time of implantation (8). In mice, relaxing the muscle by activating the beta 2 adrenergic receptor (β2AR) signaling on day 3 of pregnancy disrupts embryo movement and thus spacing, causing embryo crowding at the time of implantation (9). Outcomes of mammalian pregnancy are sensitive to adrenergic signaling regulated muscle contractions, which in turn, is indicative of how pregnancy might respond to stress levels in the mother (10). Thus, understanding how and when adrenergic muscle contractions regulate embryo movement and spacing is crucial.

Lysophosphatidic acid (LPA) signals through G protein-coupled receptors (11) and acts on the uterus through receptor LPAR3, affecting embryo spacing as well as implantation (12). A link between β2AR signaling as well as LPAR3-signaling to mediate embryo spacing has been suggested (9). However, it is intriguing to note that causing muscle relaxation by activating the β2AR signaling causes embryo overcrowding at different sites along the uterine horn, and there are often multiple embryo clusters implanted (9). Conversely, deleting LPAR3-mediated signaling in the uterus always causes embryo implantation in a single cluster either in the middle or closer to the cervical region of the uterus (11, 12). These observations suggest that although both muscle contraction and LPAR3-signaling play a role in embryo spacing, they may be targeting distinct processes in the embryo spacing pathway.

Part of the challenge in understanding preimplantation events even in a highly tractable model system such as the mouse is that the peri-implantation embryo is about 100 μm in diameter while the uterine milieu is 2-3 cm in length and 1-1.5 mm in-depth, posing a technical imaging challenge to assess both structures simultaneously. Here, using a recently developed imaging and image analysis methodology (13), we detect embryos in the mouse after entry into the uterine horn through peri-implantation stages, documenting their location as they navigate to their implantation site. We show that embryos move in two different phases displaying either clustered movement or scattering movement. We further show that while adrenergic muscle contraction is key to the initial clustered movement of the embryos, the scattering phase of embryo movement is independent of such muscle contractions but depends on the embryo as well as the LPA-LPAR3 signaling pathway. Thus, both mechanical and biological forces regulate embryo movement and spacing to achieve ideal implantation.

## Results

### Embryo movement occurs in phases: entry, unidirectional clustered movement, and bidirectional scattering movement

We used our newly developed method (materials and methods, Fig. S1) to determine embryo location at the early stages of pregnancy when embryos are present in the uterine horn, but vascular permeability has not yet begun (14). On GD3 of mouse pregnancy, the earliest evaluation of embryo location histologically has been performed at GD3 10:00h (5), and although it has been mentioned that embryo entry into the uterus occurs earlier than 10:00h (15), there has been no morphological data previously presented. We started our embryo location study at the beginning of GD3 at 00:00h. We found that on GD3 at 00:00h and 03:00h, embryos are present in clusters near the oviductal-uterine junction (100% and 71%, respectively. Fig. 1A, Movie S1, S2). We term this first phase, ‘*embryo entry*.’

**Figure 1:**
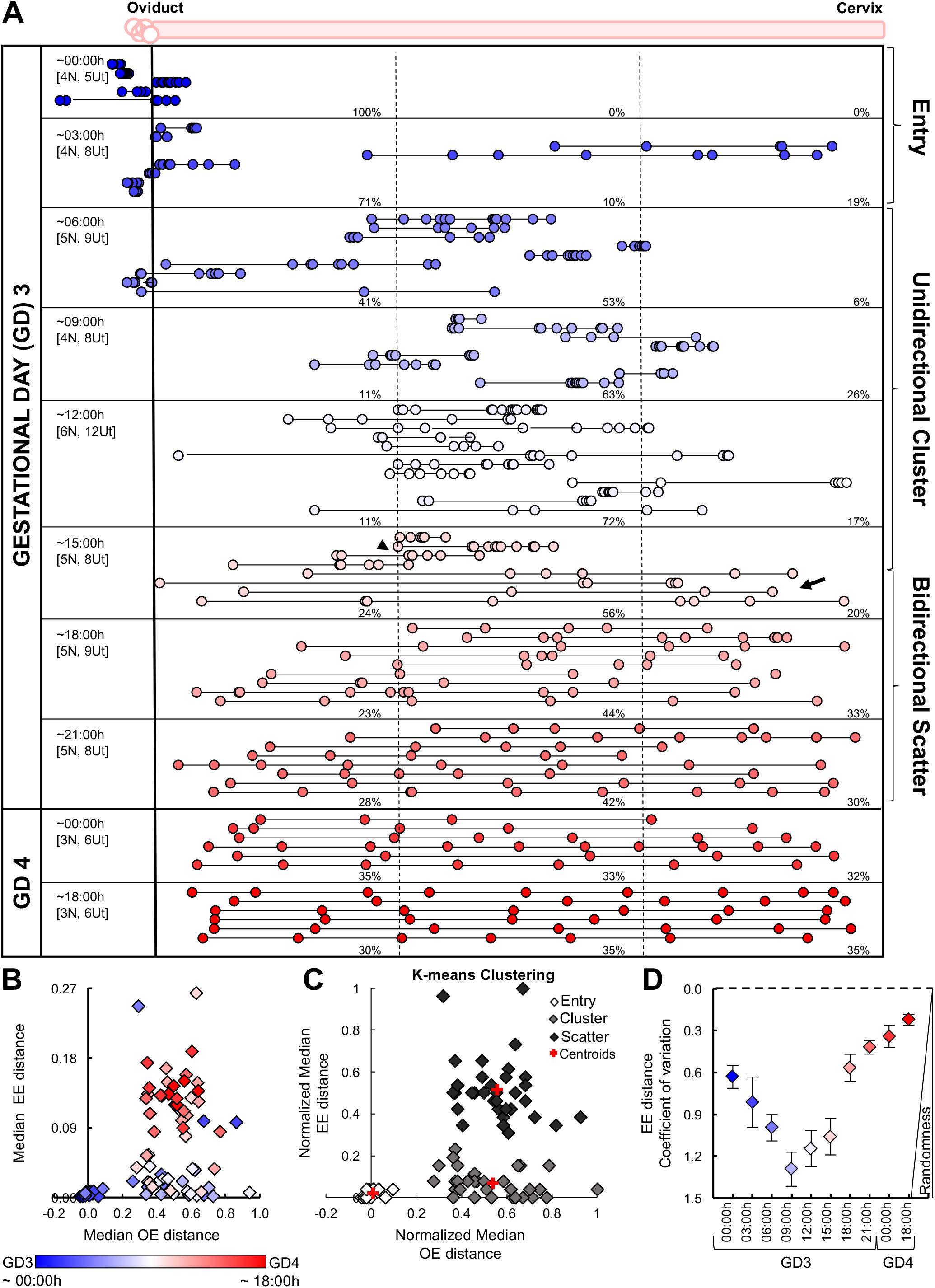
Embryo movement analysis. **(A)** Location of embryos in uterine horns at fixed time intervals on GD3 and GD4. Each circle represents an embryo, and circles connected with a line are embryos from the same uterine horn. Blue-white-red colors signify time scale where blue is the earliest time point on GD3, white is in the middle of GD3, and red is the latest time point on GD4. The left-hand column indicates the time of dissection in hours (h). ‘N’ represents the number of mice, and ‘Ut’ represents the number of uterine horns analyzed for each time point. Dotted lines divide the uterine horns into three equal segments, and percentages for each time point signify the percentage of embryos in each segment. **(B)** Classification of uterine horns from (A) based on median OE and EE distance. **(C)** k-means clustering algorithm detects three clusters corresponding to the three phases: embryo entry, unidirectional embryo clusters, and bidirectional embryo scattering. **(D)** Average Coefficient of Variation (COV) of EE distances over different time points. Time points closer to implantation and post-implantation (in red) display non-random embryo spacing distribution. *Error bars* represent S.E.M.

At GD3 06:00h, we observe that embryos remain in clusters but are present further along within the uterine horn either in the first third (41%) or in the middle fragment of the horn (53%, Fig. 1A). At subsequent time points – 09:00h and 12:00h, embryos are primarily present in clusters in the middle fragment of the horn (63% and 72%, respectively. Fig. 1A). Interestingly, at these time points, the embryos are rarely observed near the oviductal or the cervical end of the uterine horn (Fig. 1A). This movement of embryos in clusters appears to have a net unidirectional movement until embryos reach the middle of the horn. We term this second phase, ‘*unidirectional clustered movement*.‘

At GD3 15:00h, embryos are present as clusters in half of the uterine horns, (Fig. 1A, black arrowhead). In contrast, in the other half, clusters begin to separate, and embryos are present along the entire length of the uterine horn (Fig. 1A, black arrow). At 18:00h, embryos continue to disperse (Fig. 1A), and at 21:00h and beyond, fine-tuning of embryo spacing occurs along the length of the uterine horn (Fig. 1A). During this phase, embryos appear to move bidirectionally from the middle of the horn to space out evenly along the entire length of the uterine horn and eventually are equally distributed in the uterine horn. We defined this third phase as ‘*bidirectional scattering movement*.’

When mean values of OE distances are plotted as a function of time, the embryos reach a mean distance of 0.5 around GD3 09:00h (Fig. S2A). It is important to note that a mean distance of 0.5 could signify either the movement of embryos in a cluster and their arrival in the middle of the horn or could imply embryo spacing throughout the uterine horn. To distinguish between these two possibilities, it is essential to evaluate both OE and EE distances as a function of time. We observe that mean values of EE distances do not start increasing until GD3 15:00h suggesting that the embryos arrive in the middle of the horn around GD3 09:00h and stay there for ~6 hours before they begin to scatter and space (Fig. S2B).

For each horn in Fig. 1A, we plotted the median OE and EE distances against each other to visualize their relationship (Fig. 1B). We further performed the k-means clustering algorithm, and it automatically classified our data set into three groups (Fig. 1C) that correspond to the three embryo movement phases defined earlier (Fig. 1A, 1B). When measured for OE and EE parameters, horns in each of the three phases were statistically different from each other (ANOVA: OE p<0.0001; EE p<0.0001), whereas horns within the same phase were not statistically different from each other. This analysis also helped resolve outliers in the individual time points where there was a variation in embryo location likely a result of biological differences arising due to mating time (e.g., Fig. 1A, 03:00h).

### Embryo-embryo spacing approaches uniform distribution closer to implantation

Although there is variation in embryo numbers in mouse pregnancies and from one pregnancy to another in the same mouse, embryos always achieve seemingly even spacing in individual uterine horns around the time of attachment. This suggests that in the mouse, similar to the rat and rabbit, embryo spacing is not random but a guided process. To mathematically assess if embryo spacing is a non-random process, we calculated the COV of the EE distances. A larger value of COV indicates more variability in the data set in relation to the mean of the population, i.e., a more random process. As the embryo location changes through the horn at different time points on GD3, we observe differences in the COV (Fig. 1D), but as the time of attachment gets closer (GD3 21:00h, COV = 0.42), the embryo spacing distribution becomes non-random (COV closer to 0) suggesting even spacing of embryos through the horn (Fig. 1D). When mean EE distances were plotted as a function of the number of embryos for post-implantation time points on GD4 (00:00h and 18:00h), there was an inverse correlation between these two variables (Fig. S2C, R^2^=0.88). This suggests that implantation sites are not predetermined but guided by the number of embryos present in the uterine horn of the mouse.

### Unidirectional but not bidirectional phase of embryo movement relies on adrenergic smooth muscle contractions

Uterine smooth muscle contractions are thought to be a key factor for embryo movement and spacing along the uterine horn (1,4, 9, 10). If this is indeed true, then relaxing the muscle during different phases should halt embryo movement and cause overcrowding. We examined the effects of salbutamol, a muscle relaxant, by relaxing the muscle at the beginning of distinct phases of embryo movement (as determined from Fig. 1A). For disrupting muscle activity in the unidirectional clustered phase (Fig. 2A), we injected salbutamol twice on GD3 at 3:00h, and 11:00h and evaluated embryo location at 15:00h. In salbutamol-injected mice, 78% of embryos were found proximal to the oviductal-uterine junction (Fig. 2A”), as opposed to vehicle-injected mice where embryo clusters moved farther along the horn and only 45% were found proximal (Fig. 2A’). These data suggest that muscle contraction during the unidirectional phase is required for clustered embryo movement through the horn. On the other hand, vehicle (Fig. 2B’) or salbutamol (Fig. 2B”) treatment at the beginning of the bidirectional phase of embryo movement (GD3 11:00h) did not alter embryo movement patterns on GD3 at 19:00h. When median OE and EE distances were plotted we observed that muscle relaxation inhibited clustered embryo movement resulting in both lower EE and OE values in salbutamol-treated horns compared to vehicle-treated controls (Fig. 2A’”, t-test: OE p<0.0002). On the other hand, when muscle relaxation is induced prior to the scattering movement, the vehicle or salbutamol treatment does not affect OE and EE distance distribution (Fig. 2B’”, t-test: OE p<0.37). These data suggest that once the embryos arrive in the middle of the uterine horn, they initiate bidirectional movement and space out independent of the adrenergic smooth muscle contractions.

**Figure 2:**
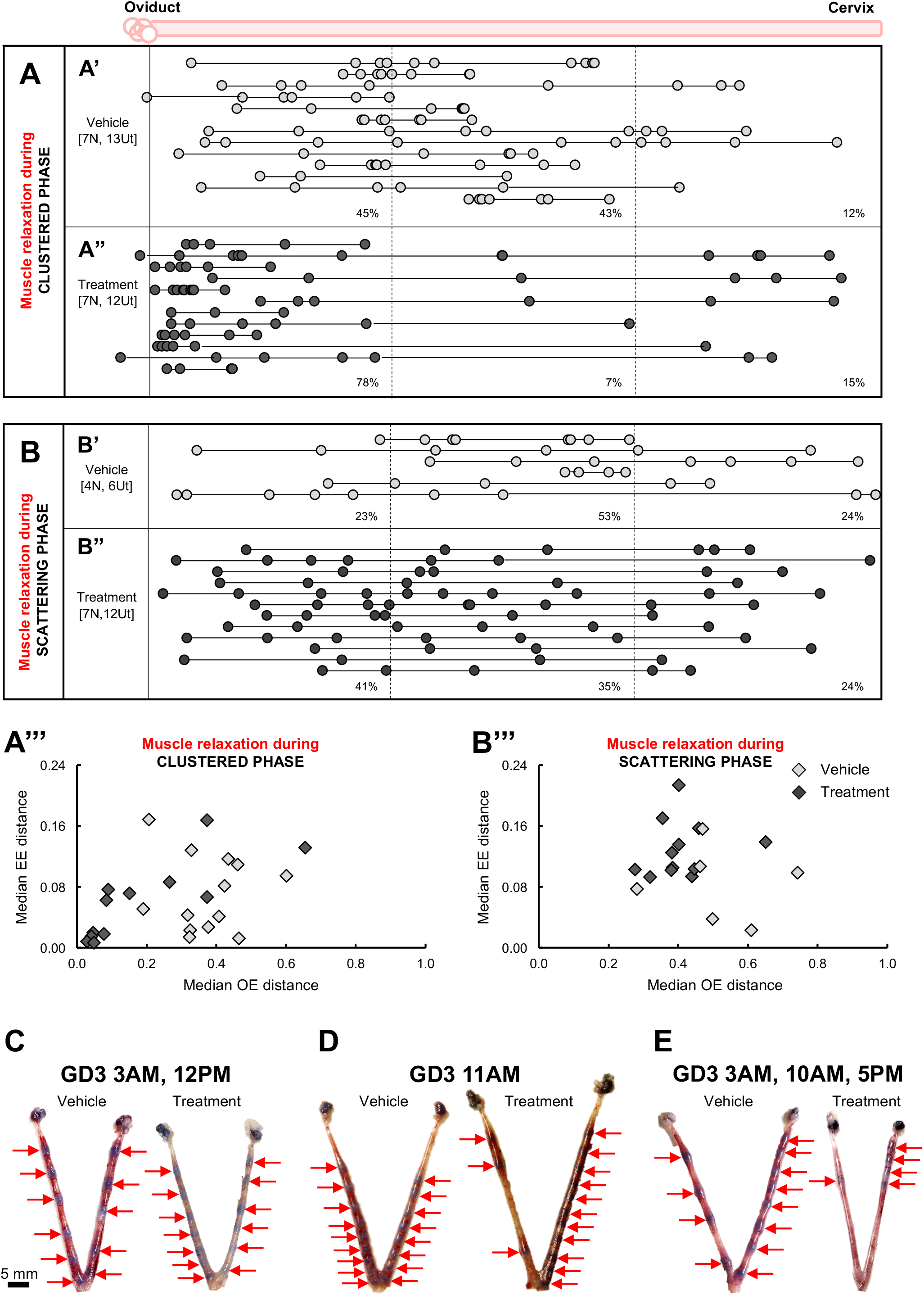
Muscle contractions regulate embryo movement in the unidirectional phase but not during the bidirectional phase. **(A)** When the uterine muscle is treated with vehicle (A’) at the beginning of the unidirectional clustered movement phase (GD3 03:00h and 10:00h), and embryo location is assessed 12 hours later (GD3 15:00h) the embryos show a distribution of 45%, 43%, and 12% in the three different uterine segments respectively. On the other hand, when the uterine muscle is relaxed with salbutamol treatment (A”), 78% of the embryos are present in the first uterine segment followed by 7% in the second fragment, and 15% are present in the third segment suggesting clustered embryo movement is reliant on muscle contractions. **(B)** If the uterine muscle is treated with vehicle (B’) or salbutamol (B”) at the beginning of the scattering movement once embryo clusters have arrived in the middle of the uterine horn (GD3 11:00h) and embryo location is assessed 8 hours later, embryo distribution is similar suggesting relaxing uterine muscle does not influence embryo scattering and spacing. (A’”) Classification of uterine horns based on median OE and EE distances for embryos in A and (B’”) for embryos in B. Analysis on GD4 18:00h to evaluate embryo location and spacing at implantation using blue dye method when: **(C)** mice are treated with vehicle or salbutamol as in (A) or **(D)** mice are treated with vehicle or salbutamol as in (B) or **(E)** when mice are treated with vehicle or salbutamol continuously starting at the beginning of the unidirectional clustered phase until a few hours before attachment (GD3, 03:00h, 10:00h and 17:00h).

To determine how long the uterine muscle contraction is required to allow for embryo scattering and even spacing for implantation, we treated pregnant mice with salbutamol twice on GD3 at 03:00h, and 11:00h as in Fig. 2A and evaluated embryo distribution with blue dye injection at GD4 18:00h (Fig. 2C, N=3). Embryo distribution and spacing are comparable to controls suggesting that if the muscle has time to recover from the effects of the muscle relaxant prior to implantation, embryo distribution and spacing are rescued, and embryos are spaced out evenly at implantation (Fig. 2C). On the other hand, if mice are treated with salbutamol thrice on GD3, at 03:00h, 10:00h, 17:00h, and evaluated for embryo distribution with blue dye injection at GD4 18:00h we find that embryos are clustered during the time of implantation (Fig. 2E, N=4). If embryos are treated with salbutamol in the clustered phase (GD3 11:00h) and assessed for implantation at GD4 18:00h with blue dye, as expected from evaluation at GD3 19:00h (Fig. 2B), embryos are spaced evenly (Fig. 2D, N=3). Thus, uterine muscle contractions allow movement of the clustered embryos, provided the muscle contracts for a sufficient amount of time prior to implantation.

### Beads are capable of movement in the unidirectional phase but not in the bidirectional phase

Embryo movement through the uterus can be due to biological communication between the embryo and the uterus, biological communication amongst the embryos, or solely caused by the embryo’s physical stimulus as an object. To distinguish between these possibilities, we injected blastocyst-sized beads in a pseudopregnant recipient uterine horn before the unidirectional clustering phase (GD2 22:00h) or before the bidirectional spacing phase (GD3 11:00h).

When beads are injected near the oviductal-uterine junction before the time of embryo entry (GD2 22:00h) and evaluated 1 hour later, beads are present as clusters with 100% of the beads in the first third of the uterine horn (Fig. 3A). When beads are injected near the oviductal-uterine junction before the time of embryo entry (GD2 22:00h) and evaluated 12 hours later, the beads are not present in clusters, but instead are spread out along the uterine length with beads present in all three segments of the uterine horn (Fig. 3B). OB and BB distance analyses show that at 1 hour post-injection, beads are in the clustered phase, whereas at 12 hours post-injection, beads are already in the scattering phase (Fig. 3E). These data indicate that in the unidirectional movement phase, beads as objects show movement along the uterine horn when injected near the oviductal end of the uterus, but the movement pattern differs from embryos as the beads fail to move in clusters over time.

**Figure 3:**
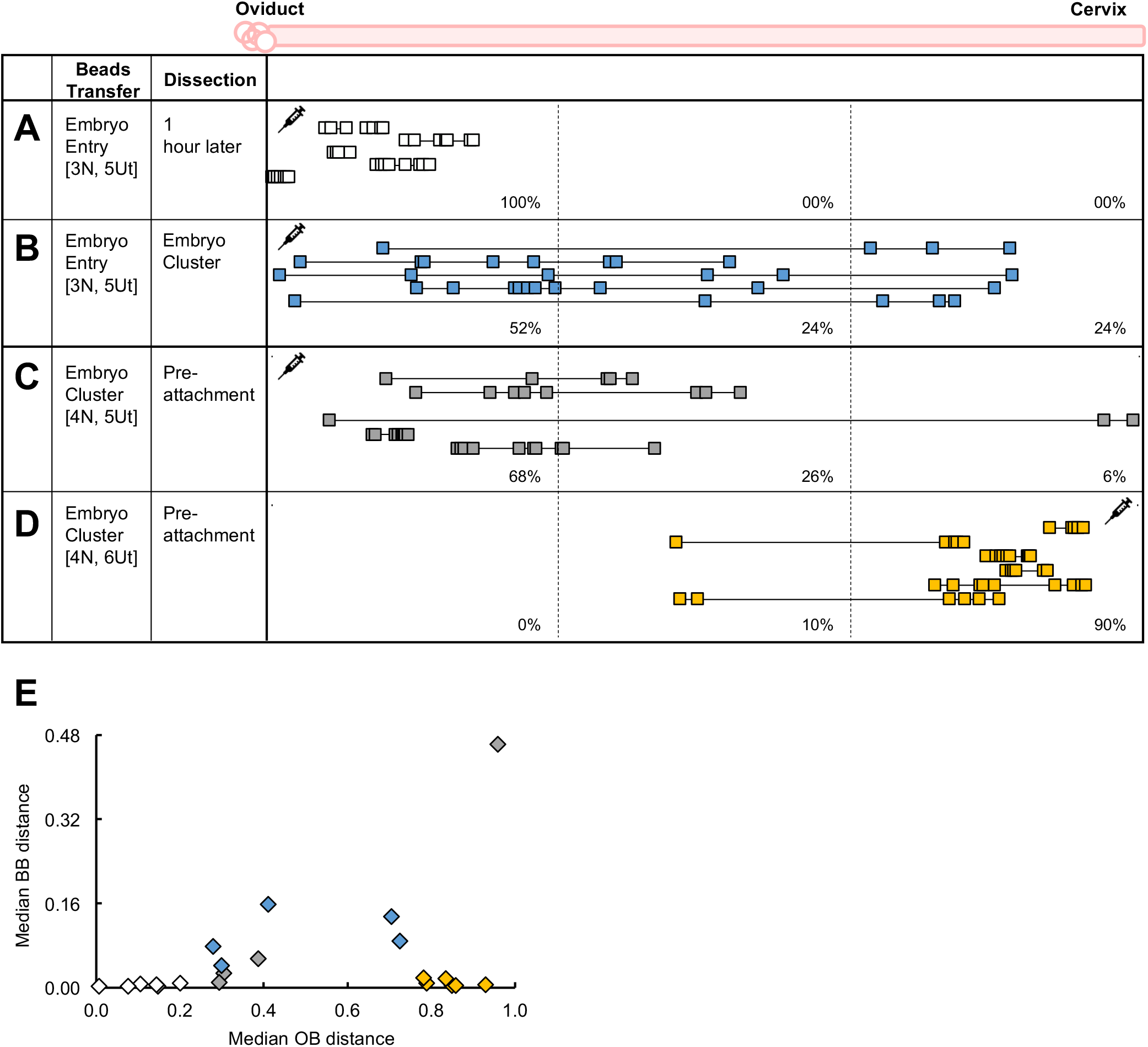
Inert beads move during the unidirectional phase but not in the bidirectional phase. Beads injected near the oviductal-uterine junctions at the time of embryo entry move in clusters 1 hour post-injection **(A)** but began scattering along the uterine horn 12 hours post-injection **(B).** Beads injected at the beginning of the scattering phase near the oviductal-uterine junction **(C)** or near the cervical region of the uterus **(D)** stay primarily near the site of injection and do not move much. **(E)** Median OB and BB distance of uterine horns in (A), (B), (C) and (D).

When beads were injected at the beginning of the bidirectional movement phase (GD3 11:00h) and evaluated 8 hours later, they stayed in clusters and failed to display much movement or spacing away from the injection site (Fig. 3C). Bead movement was sparse irrespective of whether the beads were injected near the oviductal end (Fig. 3C) or the cervical end (Fig. 3D) of the uterus, as the majority of the beads were found in the segment where the injection site was. OB and BB analysis show that beads are stuck in clusters when injected before the beginning of the scattering phase (Fig. 3E).

These results suggest that beads as physical objects move along the uterine horn in the unidirectional phase but fail to move in the bidirectional spacing phase highlighting that embryo-uterine or embryo-embryo interactions become critical to the movement of embryos in the bidirectional scattering phase.

### Movement in the bidirectional phase relies on embryo-uterine communication

LPA-LPAR3 signaling is essential for embryo spacing and on time embryo implantation (12). Embryos in *LPAR3^-/-^* uteri are often found near the cervical end upon implantation, irrespective of the embryo genotype (12). We evaluated the location of the embryos in *LPAR3^-/-^* uteri at the end of the unidirectional clustered phase (GD3 12:00h) and during the bidirectional scattering phase (GD3 18:00h and 21:00h). Embryos in the *LPAR3^-/-^* uteri are located in clusters in the second and third fragments of the uterine horn at GD3 12:00h, suggesting that they are capable of clustered unidirectional movement (Fig. 4A, 4A’ (12, 15)). On the other hand, embryos in *LPAR3^-/-^* uteri are unable to display bidirectional movement and remain in clusters in the middle of or near the cervical end of the uterine horns (Fig. 4B, 4B’ (11)). EE distances were significantly different between controls and *LPAR3^-/-^* uteri during the bidirectional scattering phase (p<0.0001) but not during the unidirectional clustered phase. These data suggest that embryo-uterine communication mediated by LPAR3, but not embryo-embryo communication plays an essential role in the bidirectional movement phase.

**Figure 4:**
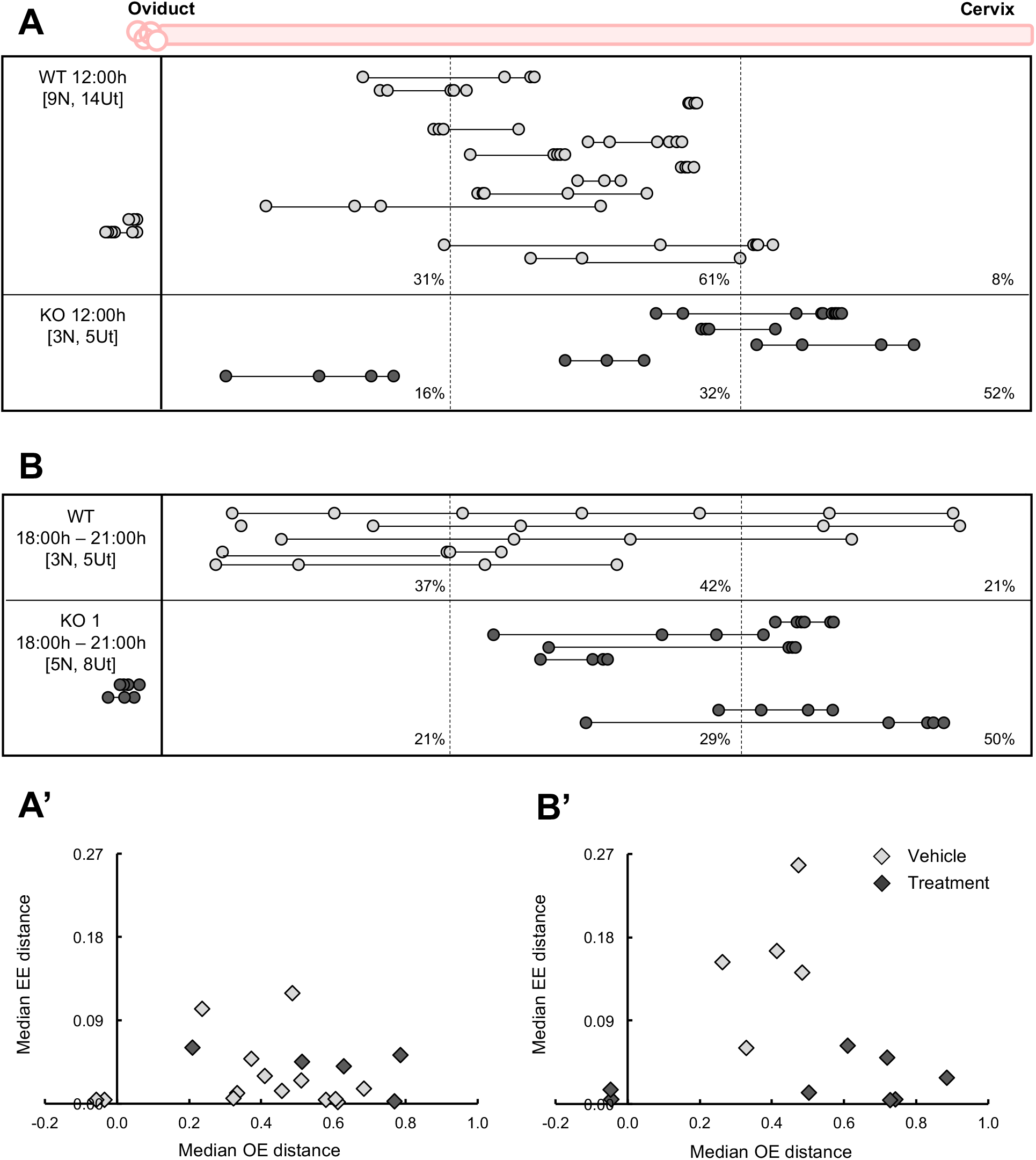
LPAR3 dependent embryo-uterine communication is essential for the bidirectional scattering of embryos. **(A)** Embryos are present in clusters in the middle and cervical fragments of the control and *LPAR3^-/-^* uteri at the end of the unidirectional clustered movement phase (GD3 12:00h). **(B)** While embryos in the control horns scatter between GD3 18:00h - 21:00h, embryos in *LPAR3^-/-^* horns are stuck near the cervix in clusters.

## Discussion

Our studies reflect how using 3D imaging to capture the uterine structure in its entirety as well as identifying embryos, and their location allows for a detailed quantitative analysis of early events in mammalian pregnancy. Previous work in the mouse on embryo spacing has been performed in either whole mount at a time when the implantation sites or embryonic deciduae occupy a substantial portion of the uterine horn (16) or as 2D histological sections near the time of implantation (5). In order to make accurate estimates of even and uneven spacing, embryo distribution analysis needs to be performed when the object to be distributed (the embryo) occupies less than 5% of the area it will be distributed in (the uterine horn) (2). Thus, we applied our methodology to carry out quantitative embryo location analysis in the mouse during periimplantation events.

### Embryo location and movement: similarities and differences between mammalian species

Previous work with model organisms of mammalian pregnancy, such as the rat and the rabbit have described quantitatively in detail, events of early embryo movement through the uterine tissue. We find differences and similarities in our data collected in mice in comparison to the data with rats and rabbits.

#### Embryo Entry

We observed time points where embryos are in the oviduct as well as the uterine horn, and within the next 3 hours, find all the embryos as a cluster near the oviductal-uterine junction inside the uterine horn. This suggests that similar to the rabbit (2), mouse embryos enter rather quickly as a cluster, not one by one.

#### Movement through the horn

In rats, irrespective of the time point analyzed, embryos are always found in the first segment of the uterine horn, and in the rabbit, blastocyst progression is continuous, based on the mean/median embryo location and the progressive increase in the EE distance. This suggests that embryos in the rat and the rabbit uterus move unidirectionally, and scattering begins as the embryos are moving and spacing unidirectionally. These data are in contrast to our data with the mouse as there are time points where embryos are sparsely found in the first third of the uterine horn, and EE distances stay small in the unidirectional phase and only start to increase in the bidirectional scattering phase of embryo movement.

Although there seems to be an absence of clustered movement in the rabbit, Boving did note that there is a separation between blastocyst location and the end zones of the uterus, implying blastocysts are not present near the oviductal or cervical end of the uterus (2). This is similar to the end of the clustered phase movement in the mouse, where embryos are close to each other but far off from either end of the uterus in the mouse. Evolutionarily, why there is a time when both ends of the uterus in the rabbit, as well as the mouse, does not contain embryos is an intriguing observation and warrants further investigation.

In contrast to previous data with the mouse (5), where embryo spacing is suggested to begin at 10:00h, our data clearly shows that embryos stay clustered and start spacing around 15:00h on GD3.

#### Spacing and Implantation

While there are differences in how embryos move through the uterine horn and the sequence in which they achieve spacing, rats (3, 4), rabbits (2), and mice (this study) all show even embryo spacing at the time of implantation as determined by the COV of the EE distances. This, along with the fact that uterine horns vary in length, and EE distance postimplantation is inversely proportional to the number of embryos in the uterine horn, supports the idea that implantation sites are not predetermined but are formed once embryos enter the horn and begin interacting with the uterine environment.

### Adrenergic muscle contraction is responsible for embryo movement but not embryo spacing

We determined that adrenergic uterine contractions are required for the unidirectional movement of embryo clusters but not the bidirectional movement for embryo scattering. Overcrowding near the oviductal-uterine junction was observed when the muscle is relaxed at the time of embryo entry and evaluated prior to attachment. A similar overcrowding effect was previously reported with salbutamol in mice (9) and with relaxin in rats (4). Although interfering with contractile activity prior to embryo arrival at the center of the horn causes overcrowding of embryos and compromises pregnancy outcomes ((9) and our study), it is key to note that overcrowding of embryos at implantation depends on the amount of recovery time between removal of the muscle relaxant stimulus and embryo attachment. A 13-hour gap between the last injection of salbutamol (GD3 11:00h) and embryo attachment (GD3 24:00h) allows for recovery of embryo movement once the effect of the muscle relaxant wears off and spacing is observed at GD4. On the other hand, a 7hour gap between the last injection of salbutamol (GD3 17:00h) and embryo attachment causes overcrowding of embryos at GD4, suggesting a lack of sufficient recovery time between removal of the muscle relaxant and embryo attachment. These data are similar to the rat study, where if rats were treated with relaxin post embryo entry but were allowed a recovery period of ~24 hours before attachment (4), the embryos were found to be distributed evenly throughout the horn but if embryos are continuously exposed to relaxin after embryo entry until the time of attachment (8), embryo overcrowding near the oviduct is observed. We speculate that once the embryos are positioned in the center of the horn in the mouse and the uterus has presumably counted the embryos, the adrenergic uterine contractions do not play a role in embryo spacing.

### Inert objects display movement in the unidirectional clustered phase but fail to do so in the bidirectional scattering phase

Beads as inert objects move in the uterine horn only in the first phase of movement and not during the second phase. Further, bead movement begins as clusters, but over 12 hours, the beads disperse and break out of clusters suggesting continuing movement in clusters could be a function of embryo-embryo or embryo-uterine interactions. Bead movement in the unidirectional phase, along with our data that muscle contraction is required for the movement of embryos during the unidirectional phase, suggests that this phase of object movement is characterized by passive movement of physical objects under the regulation of uterine peristalsis. This passive movement of objects is time-sensitive as beads do not move when introduced into the uterine horn during the second phase of movement. Uterine contractile activity by itself was not enough to move beads during the scattering phase. Thus, the second phase of movement relies on active communication between the embryo and the uterus to facilitate even spacing throughout the horn.

### LPA-LPAR3 signaling plays a role in the bidirectional scattering phase of embryo movement

While an embryo is key to even spacing, studies with genetic mutants where control embryos are incapable of spacing in gene-deleted uteri (e.g., LPAR3 (12), cPLA2 (17)) suggest that there is an active embryo-uterine component to embryo spacing. Because *LPAR3* mRNA was downregulated in the salbutamol treated uteri (9), but patterns of overcrowding are distinct between *LPAR3^-/-^* mice (embryo crowding in a single cluster, near the cervix), and salbutamol treated mice (embryo crowding in multiple clusters at different sites along the uterine horn) we wanted to determine the effects of LPAR3 deletion on embryo distribution and location. Our study indicates that *LPAR3^-/-^* uteri behave similarly to control uteri during the unidirectional clustered movement, allowing the embryos to travel through the uterine horn in the first phase of movement and arrive near the middle or the cervical portion of the uterus. Still, these uteri cannot respond to the embryos to initiate the bidirectional scattering movement. The defect in embryo spacing in *LPAR3^-/-^* uteri is attributed to the lack of a specific subset of contraction, which can otherwise be induced in a control uterus by an LPAR3-specific agonist (15). Thus, embryo-uterine communication mediated by LPAR3 and contractions that lie downstream of this communication are crucial to embryo spacing.

Boving has implicated that muscle activity that propels embryo movement is distinguished by either spontaneous contractions or stimulated contractions. Spontaneous contractions can be induced by prostaglandins (such as PGF2α) and are suppressed by estrogen and relaxin (18). Progesterone, on the other hand, acts on uterine muscle by conditioning it and reducing the spread of contractile activity (19). Stimulated contractions are likely induced by the embryo itself. We speculate that the first phase of the unidirectional clustered movement is under the influence of spontaneous contractions that involve the adrenergic signaling pathway. These contractions are likely inhibited at the time of the nidatory peak of estrogen (20, 21), and movement of embryos is then guided by embryo stimulated contractility of the uterus. These second set of contractions are likely under the regulation of ovarian hormone progesterone as well as LPAR3, and these stimulated contractions ensure equal spacing of embryos prior to implantation.

Our study provides a deeper understanding of the embryo movement process that could be dependent on physical forces (muscle contractions) or biological forces (the embryo or LPA-LPAR3 signaling). This understanding is essential because modulating these processes will be key to manipulating early events in implantation for pregnancy success and for development of novel methods of contraception.

## Materials And Methods

### Animals

All animal research was carried out under the guidelines of the Michigan State University Institutional Animal Care and Use Committee. Crl: CD1(ICR), wild-type C57BL/6, and *Lpar3^tm1JCh^* (*LPAR3^-/-^*) mice (12), aged 6 to 8 weeks, were maintained on a 12 hours light/dark cycle. Adult females were mated with fertile or vasectomized wild-type males to induce pregnancy or pseudopregnancy. The appearance of a vaginal plug was identified as the gestational day (GD) 0.5. For CD1 females, uterine dissections were performed at 3-hour intervals, starting from midnight on GD3 at 00:00 hours (h) and on GD4 at 18:00h. *LPAR3^-/-^* and C57BL/6 mouse uterine dissections were performed at 12:00h or between 18:00h - 21:00h on GD3. For detecting implantation sites on GD4, 200 μl of 0.5% Evans blue dye (MP Biomedicals, ICN15110805) in phosphate-buffered saline (PBS) was injected into the lateral tail vein of the pregnant mouse 15 minutes prior to sacrificing the mouse. Uteri were then photographed in white light to observe implantation sites (14).

### Beads transfer

Spherical agarose beads in PBS (4-10 beads/horn; Affi-Gel Blue Media, 153-7302) with 75 −150 μm in diameter were injected into the uterine horns between 20:00h and 24:00h on GD2 of pseudopregnancy, or at 11:00h on GD3 of pseudopregnancy. Beads were injected either near the oviductal-uterine junction region or the region of the uterus closest to the cervix. Mice were sacrificed on GD3 at 12:00h or 21:00h.

### Drug T reatments

Salbutamol (Alfa Aesar, A18544) was dissolved in a 10% ethanol solution made in PBS and injected intraperitoneally at 2 mg/mouse (9). Mice in the vehicle group received injections of a 10% ethanol in PBS solution. Vehicle or salbutamol was administered on GD3 either once at 11:00h, twice at 3:00h and 11:00h, or thrice at GD3 3:00h, 10:00h and 17:00h.

### Whole-mount immunofluorescence

Whole-mount immunofluorescence was performed as described previously (13). Briefly, uteri were dissected and fixed in DMSO:Methanol (1:4), rehydrated for 15 minutes in 1:1, Methanol: PBST (PBS, 1 % Triton X-100) solution, followed by a 15 minutes wash in 100% PBST solution. Samples were incubated in a blocking solution (PBS, 1% Triton X-100, 2% powdered milk) for 2 hours at room temperature. Uteri were incubated with primary antibodies diluted in blocking solution (1:500) for five nights at 4°C. Subsequently, the samples were washed six times for 30 minutes each using PBST and then incubated with secondary antibodies diluted in PBST (1:500) for two nights at 4°C. The uteri were then washed six times for 30 minutes each using PBST, followed by a 30 minutes dehydration in 100% methanol, an overnight incubation in 3% H_2_O_2_ solution diluted in methanol, and a final dehydration step for 30 minutes in 100% methanol. Samples were cleared using a 1:2 mixture of Benzyl Alcohol: Benzyl Benzoate (Sigma 108006, B6630).

Primary Antibody used was: ECAD (M106, Takara Biosciences). Secondary antibodies conjugated Alexa Flour IgGs were obtained from Invitrogen, and Hoechst (Sigma Aldrich, B2261) was used to stain the nucleus.

### Confocal microscopy

Uteri were imaged using a Leica TCS SP8 X Confocal Laser Scanning Microscope System with white-light laser, using a 10x air objective. For each uterine horn, z-stacks were generated with a 7.0 μm increment, and tiled scans were set up to image the entire length and depth of the uterine horn (13). Images were merged using Leica software LASX version 3.5.5 (Fig. S1A).

### Image analysis for embryo location

Commercial software Imaris v9.2.1 (Bitplane) was used for image analysis. The confocal LIF files were imported into the Surpass mode of Imaris. For embryo location analysis structures of the oviductal-uterine junction (green arrow), embryos (white arrows), beads, and horns were created as 3D renderings using the Surface module (Fig. S1A, A’, A”). With the Measurements module, the three-dimensional Cartesian coordinates of each surface’s center were identified and stored. The Cartesian coordinates of the orthogonal projection onto the XY plane were used to calculate the distance between the oviductal-uterine junction and an embryo (OE), the distance between the oviductal-uterine junction and a bead (OB), the distance between adjacent embryos (EE), the distance between adjacent beads (BB), and the horn length. All distances were normalized to the length of the horn to compensate for uterine horn length differences amongst mice unless otherwise specified. Horns with less than three embryos/beads were excluded from the analysis. Finally, these distances were used to map the location of the embryos/beads relative to the length of the uterine horn. To confirm the robustness of our embryo location method, we compared the results of our embryo location analysis to the established blue dye injection method at the time of embryo implantation (14). When we compare our embryo location data at GD4 18:00h, it overlaps with the data generated using a blue dye permeability assessment (Fig. S1B) and also gives us the precise location of the embryo in the uterus.

To further assess the relative location of the embryo in the horn, we divided the uterine horn into three equally spaced segments – segment closest to the oviduct, middle segment, and a segment closest to the cervix (dotted lines in Fig. S1B) and assessed the percentage of embryos present in each section. Embryos present in the oviductal region close to the oviductal-uterine junction were accounted for in the first segment. These quantitative measurements are useful for comparing our data to the embryo location generated in the rabbit and the rat (2, 4, 22).

### Coefficient of Variation

We computed the coefficient of variation (COV) as the standard deviation of EE distances divided by the mean of the EE distances for each horn of the time points assessed (2). The mean value of all COVs for all horns at a particular time point was plotted as a function of time.

### K-means clustering

The relationship between OE and EE median distances was analyzed using the k-means clustering algorithm implemented on MATLAB. This method aims to divide *n* data points into *k* clusters in which each data point belongs to the cluster with the nearest mean (cluster centroid). As a preprocessing step for k-means analysis, each data set was normalized to a range −1 to 1 (23).

### Statistical Analysis

Statistical analyses were performed with Graph Pad Prism. ANOVA was used to statistically analyze OE and EE distances amongst uterine horns and different time points. To compare the vehicle and the treatment for muscle contraction analysis, and the *LPAR3^-/-^* with controls, the unpaired two-tailed t-test was performed with Welch’s correction.

## Acknowledgments

We thank Sarah Fitch, Anna Coronel, and Devan Patel for assistance with data collection. We thank Prof. Jerold Chun for the *LPAR3^-/-^* mice. We thank Dr. Amy Ralston, Dr. Asgerally Fazleabas, and Dr. Jorge Barreda for critical feedback and suggestions on the manuscript.

---

**Supplementary Figure 1:**
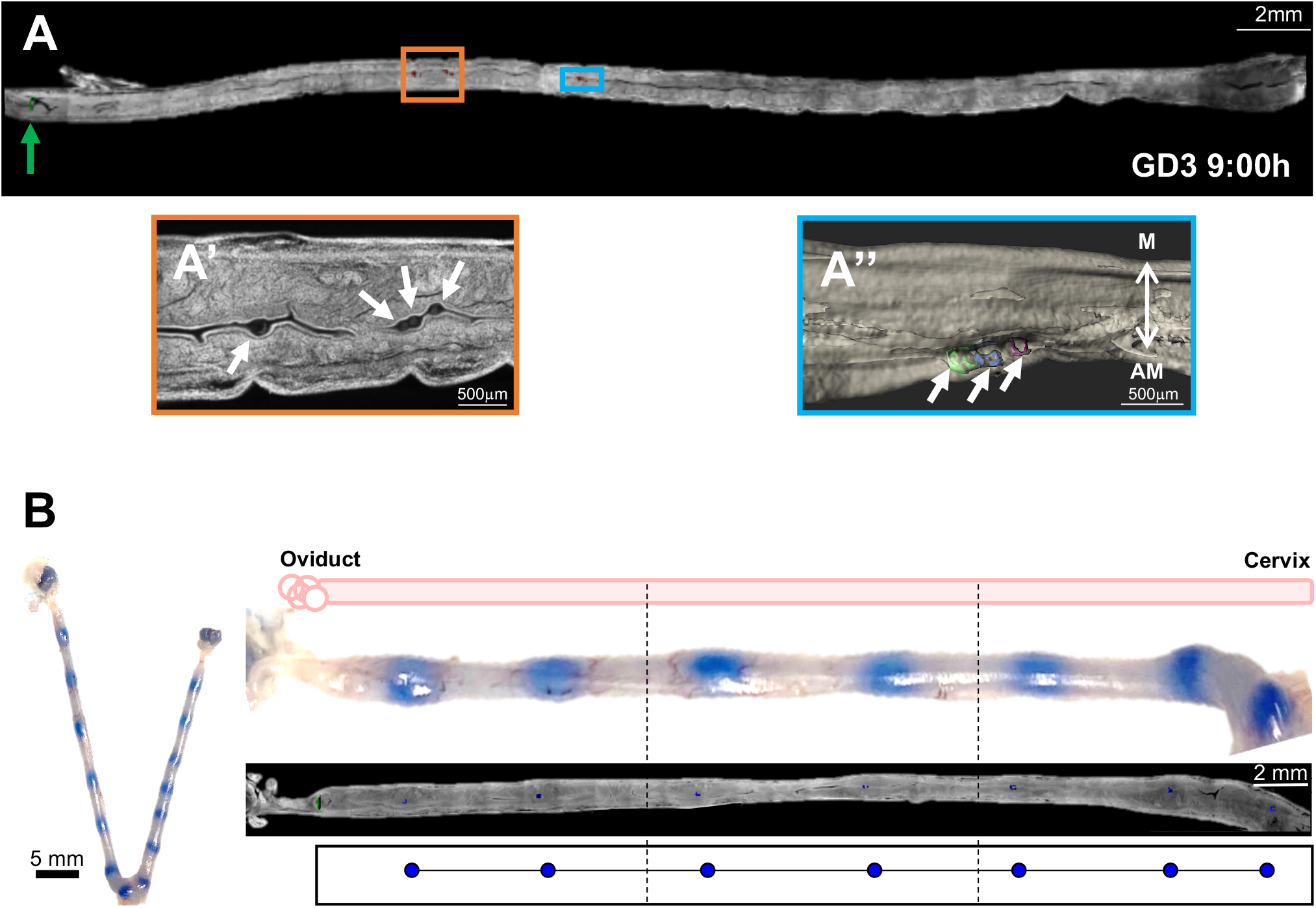
Methodology to measure embryo location along the uterine horn. **(A)** 2D optical slice of a GD3, 09:00h uterus stained with Hoechst (grey). The green arrow points to the oviductal-uterine junction. **(A’)** Magnified region for the orange rectangle in (A) showing embryos in a cluster (white arrows) on the 2D optical slice in A. **(A”)** Magnified region for the blue rectangle in (A) showing 3D reconstruction of the uterine lumen (grey) and embryo surfaces along the uterine horn (blue, green, pink surfaces). **(B)** Uterine implantation sites at GD4 18:00h shown with blue dye injections. Embryo location analysis using the methodology described in the methods. Blue circles represent embryo location with our described methodology showing similarities between the imaging-based embryo location method and the blue dye permeability method.

**Supplementary Figure 2:**
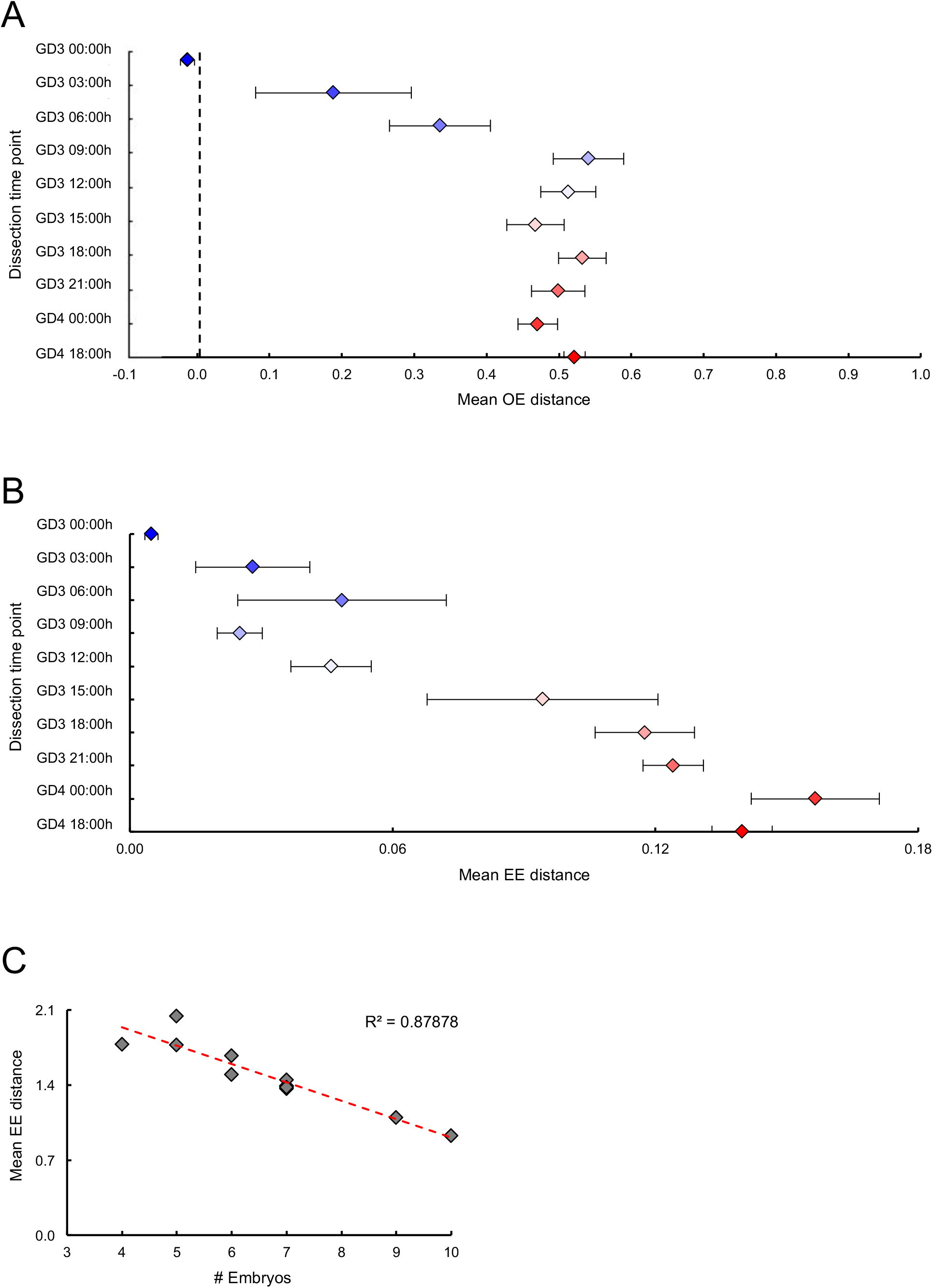
EE and OE distances plotted for each time point analyzed for the time course. **(A)** Mean EE and **(B)** Mean OE distances plotted over the time course of embryo movement. **(C)** An inverse correlation was observed between the number of embryos and EE distance for postimplantation time points (GD4 at 00:00h and 18:00h).

**Movie S1** (separate file). Movie displaying a 3D extended view of 2D optical planes of a GD3 00:00h uterus followed by a zoom into the oviductal-uterine junction and individual slice view visualization. E-CAD staining marks oviductal and uterine epithelium, and Hoechst stains all cell types in the full-length uterine horn with the ovary on the left and the cervical end on the right. Embryos, as stained with Hoechst, are present in the oviductal end.

**Movie S2** (separate file). Movie displaying staining of a GD3 03:00h uterus near the oviductal-uterine junction. E-CAD staining marks oviductal and uterine epithelium. Embryos, as stained with Hoechst, are present in the uterine opening of the junction.

